# In silico Analysis of Transcriptomic Profiling and Affected Biological pathways in Multiple Sclerosis

**DOI:** 10.1101/2021.08.15.456398

**Authors:** Rutvi Vaja, Harpreet Kaur, Mohit Mazumder, Elia Brodsky

**Affiliations:** School of Science, Navrachana University, Gujarat, India; Pine Biotech, New Orleans, USA

**Author notes:** **Emails of Authors:** Rutvi Vaja; Harpreet Kaur; Elia Brodsky; Mohit Mazumder. **Correspondence:** Rutvi Vaja, School of Science, Department of Biomedical Sciences, Navrachana University, Vadodara, Gujarat, India.

**Keywords:** Multiple sclerosis, Huntington’s disease, Myelinated Axons, Grey Matter, White Matter

## Abstract

Multiple sclerosis (MS) is a chronic autoimmune, inflammatory neurological disease that is widely associated with Grey and white matter degradation due to the demyelination of axons. Thus exposing the underlying causes of this condition can lead to a novel treatment approach for Multiple Sclerosis. The total RNA microarray processed data from GEO for Multiple sclerotic patients was comprehensively analyzed to find out underlying differences between Grey Matter lesions (GML), Normal appearing Grey Matter (NAGM), and Control Grey matter at the transcriptomics level. Thus, in the current study, we performed various bioinformatics analyses on transcriptional profiles of 184 samples including 105 NAGM, 37 GML, and 42 Controls obtained from the NCBI-Bio project (PRJNA543111). First, exploratory data analysis based on gene expression data using principal component analysis (PCA) depicted distinct patterns between GML and CG samples. Subsequently, the Welch’s T-test differential gene expression analysis identified 1525 significantly differentially expressed genes (p.adj value <0.05, Fold change(>=+/-1.5) between these conditions. This study reveals the genes like *CREB3L2*, *KIF5B*, *WIPI1*, *EP300*, *NDUFA1*, *ATG101*, *AND TAF4* as the key features that may substantially contribute to loss of cognitive functions in Multiple sclerosis and several other neurodegenerative disorders. Further, this study also proposes genes associated with Huntington’s disease in Multiple sclerotic patients. Eventually, the results presented here reveal new insights into MS and how it affects the development of male primary sexual characteristics.

## Introduction

As per the Multiple sclerosis association of America (MSAA), Multiple sclerosis(MS) is an auto-immune, demyelinating disease that mainly affects the central nervous system (1). The main symptom of MS is the demyelination of axons. MS is the most common cause of neurological disability in young children (2). It is a long-term disease that affects the brain, spinal cord, and optic nerves (3). There are 4 subtypes of MS which are categorized based on relapses, attacks, and exacerbations (1). The types of MS are namely: Clinically isolated syndrome (CIS), Relapsing-remitting MS (RRMS), Secondary-progressive MS (SPMS), and primary progressive MS (PPMS) (1). Among all these sub-types RRMS occurs in 80-85% of patients, which is the most common form and involves episodes of increasing symptoms followed by periods of remission (1). Over time then RRMS proliferates and progresses to SPMS and PPMS stages of MS (4). As per the National MS Society, there are now 2.8 million people worldwide with reported cases of MS (5). MS is expressed differently in different phenotypes. MS and its subtypes have a highly diverse biological and clinical background for each patient depending upon the stage of MS showcased. MS has been originally regarded as an auto-immune disease affecting white matter (6). Recently histopathological studies have shown that MS also heavily affects the Grey matter and its cortical regions (6). Damage to grey matter starts at the early onset of the MS disease which further affects the cognitive functions of the brain (6). The causes of grey matter damage are unclear, at the present.

In 1962, Brown Wells and Hughes described Grey matter lesions(GML) as a symptom of MS in selected MS autopsy samples (7). The authors used conventional immunohistochemistry and detected a total of 1,595 lesions in 22 MS cases, of which 26% were located in the cortical GM or at the border of the WM and GM (8). Grey matter lesions are classified into 4 subtypes (Type 1, 2, 3, and 4)based on the location they develop on (7). Among the type 4 lesions occur in the cortex affecting all six layers (9, 10). Lesions affecting the cortical areas of the brain are designated as either Type III or Type IV lesions (11,12,13). Many of the recent post-mortem studies have shown that Grey matter lesions are found in the thalamus and Caudate nucleus (14). Other parts where GML occurs are namely putamen, pallidum, claustrum, amygdala, hypothalamus, and substantia nigra (14). Early involvement of GML is found in all MS phenotypes and all MS subtypes as well (15). Both GM lesions number and GML volume increase and progress over a given period (16). The number of GML in primary cortical areas of the brain directly affects the motor functions and causes motor dysfunction (17). Overall GML is associated with cognitive impairments and cognitive dysfunction (18). Whereas GML about specific areas like cortical or hippocampal is associated with impaired visuospatial memory and processing speed (19,20).

The main goal of our study was to identify the underlying reasons at the transcriptomics level for Grey Matter degeneration in Multiple sclerosis which is originally a White matter degenerative disease. Our main concern was to investigate the affected biological pathways due to the formation of lesions in Grey Matter after diagnosis of Multiple sclerosis. We wanted to find out defects at the transcriptional level to better understand the proliferation, formation, and progression of lesions in the Grey Matter. As the GML expression is tissuespecific depending upon the phenotypes in which they occur, finding the underlying genes that cause lesions, Cognitive function loss and inflammation in MS could be very helpful for future treatments. It is to be noted that, patients with MS lesions showed larger cognitive disabilities than normal MS patients without lesions (21).

Hence, in the present study, we investigated transcriptomics profiles of Normal appearing grey matter (NAGM) and Grey matter lesions (GML) and healthy control samples (CG) to understand the affected Biological Pathways and genes behind the appearance of GML using various bioinformatics techniques. After identifying the significant genes between these groups; we scrutinized the affected biological pathways as the disease progresses to Normal appearing grey matter (NAGM) and Grey matter lesions (GML) samples.

## METHODS

### Data sets

In this study, processed microarray data was taken from the NCBI GEO [GSE131282] The dataset of this project was generated by Florian Geier et Al.(2020) and published as a bio project on NCBI with accession number PRJNA543111. This was our discovery data set. Transcriptome data contains microarray processed quantile normalized values using the Bioconductor package Lumi (version 2.32.0) of total RNA extracted from grey matter tissue from the Illumina HumanHT-12 V4.0 expression bead chip platform. The data set of the following Bio project is shown in Table 1 below. The validation dataset of this project was generated by Florian Geier et Al.(2020) and published as a bio project on NCBI with accession number PRJNA543111. The GEO Accession number was GSE131279. Transcriptome microarray data contain quantile normalized values of RNA expression extracted from grey matter tissue using the Illumina HumanHT-12 V4.0 expression bead chip platform. This data set had the following number of samples shown in the table 1 below.

**Table 1.** Datasets used in the study.

| Discovery Data (Main Dataset) |  | Validation Dataset |  |
| --- | --- | --- | --- |
| Disease State | Number of Samples | Disease State | Number of samples |
| Normal Appearing Grey matter | 105 | Normal appearing Grey Matter | 64 |
| Grey Matter Lesions | 37 | Control samples | 42 |
| Control Grey Matter<br>(Healthy) | 42 |  |  |

### Data Pre-Processing

The microarray processed data was quantile normalized signal data. The processed data had Illumina probe ids. We mapped probe IDs with the gene symbols using the SOFT family files.

### Exploratory Analysis

Exploratory analysis facilitated the examination of variation between all samples, including healthy and diseased patients samples (Normal appearing grey matter and Grey matter lesions). To understand the patterns in the data exploratory data analysis was performed using the Principal component analysis module integrated on the T-bio info server (https://server.t-bio.info/). PCA is a dimensionality reduction technique that discerns the variability between the samples (22). PCA was performed in four independent conditions: (1) All 3 conditions (Grey matter lesions Vs Normal appearing Grey Matter Vs Healthy Control Samples); (2)Grey matter lesions Vs Normal appearing Grey Matter; (3) Grey matter lesions Vs Control Grey Matter; and (4) Normal appearing Grey matter Vs Control Grey Matter. Eventually, the PCA scatter plots were used to determine the patterns.

**Figure 1:**
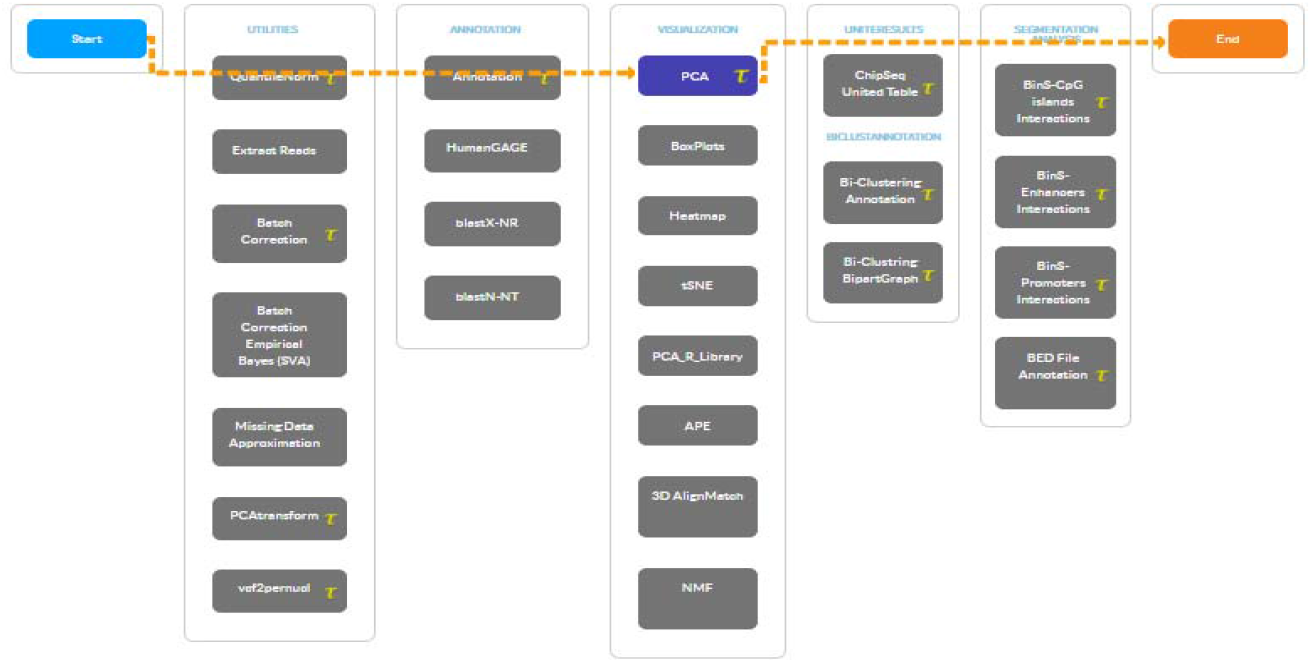
Screenshot of PCA Pipeline on T-bioinfo Server.

### Differential Gene expression Analysis

Differential Gene Expression (DGE) analysis was conducted by contrasting Grey matter lesions with Control grey matter samples. The differential gene expression (DGE) analysis was performed using the Welch’s T-test (23). Welch’s T-test is a modification of statistical analysis used for getting statistical significance for unequal variances (23). When two groups have unequal sample sizes and variances –a Welch’s Test can be applied (24). The Welch’s Test uses a one-step approach for DGE analysis and combines several methods. The significant genes were identified with the threshold of (p.adj value <0.05, Fold change (>= ±1.5) (39).

### Assessment of Discriminatory potential of Significant genes

Next, to assess and visualize the potential of significant genes in distinguishing both the classes of samples PCA and H-Clustering were performed with the selected set of significant RNA genes only. H-clustering (Distance: Euclidean, Linkage: average) was performed to understand the discriminatory potential of significant genes in distinguishing the GML and Control samples based on their gene expression. Eventually, heatmaps were drawn to visualize the gene expression patterns among different classes of samples.

### Gene Enrichment Analysis

To delineate the biological implication of the significant genes, the gene enrichment analysis for Gene Ontology (GO) terms was performed using the annotation module integrated on the T-bio info server. Further, significantly (p.adj value <0.05, Fold change(>= ±1.5) enriched KEGG pathways were identified using the Enrichr (25). Enrichr is an easy-to-use intuitive enrichment analysis web-based tool providing various types of visualization summaries of collective functions of gene lists (25). Notably, enriched pathways and GO terms were also identified using the Enrichr Platform for significant up-regulated and down-regulated genes. Further, the Enrichr was also used to identify the associated Gene Ontology terms and biological pathways with significant up-regulated and down-regulated genes.

## RESULTS

In the current study, we explored transcriptomics data of Normal appearing grey matter (NAGM), Grey matter lesions (GML), and healthy control samples using various insilco techniques to delineate the underlying gene signature and biological pathways. The complete workflow of the study is represented below in Figure 2.

**Figure 2:**
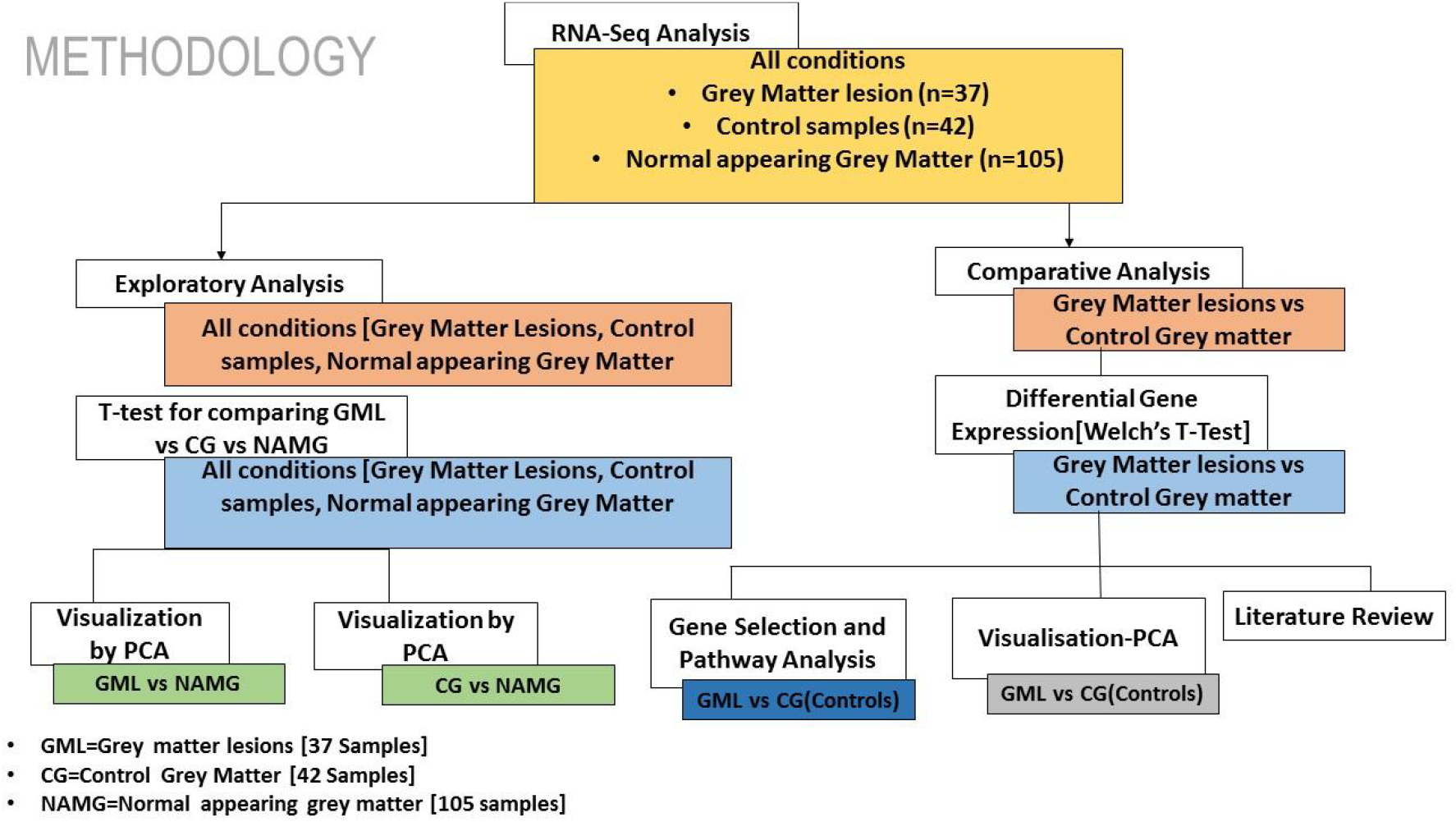
Work Flow of the study representing the key steps.

### Exploratory Data Analysis

Data were explored by using principal component analysis (PCA). We did PCA to understand the variation between the 3 groups, namely Normal appearing Grey Matter, Grey Matter lesions, and Controls. Figure 3A shows the variation between GML and NAGM. The PC1 is 84.53% along with PC2 as 11.54% and PC3=0.60%. It is evident from the PCA of this exploratory analysis that there is something that triggers the variation between GML and NAGM. Notably, variation between NAGM and Control samples is very diverse where PC1 as 81.94%, PC2 as 14.09% and PC3=0.70% as shown in Figure 3B. Figure 3C represents variation between Control Grey Matter and Grey Matter lesion samples. The variation between 3 PCAs is given as PC1=69.69%, PC2=27.31% and PC3=0.54%.Further, NAGM and CG are not separating from each other based on PCA results, here PC, PC2 and PC3 represent 82.39%, 13.74% and 0.48% variation of the data, respectively as shown in Figure 3D. Since the main focus of our study was to find out the underlying reason for Grey matter lesions and their degeneration, we decided to go with the Variance between GML and CG to further our analysis.

**Figure 3:**
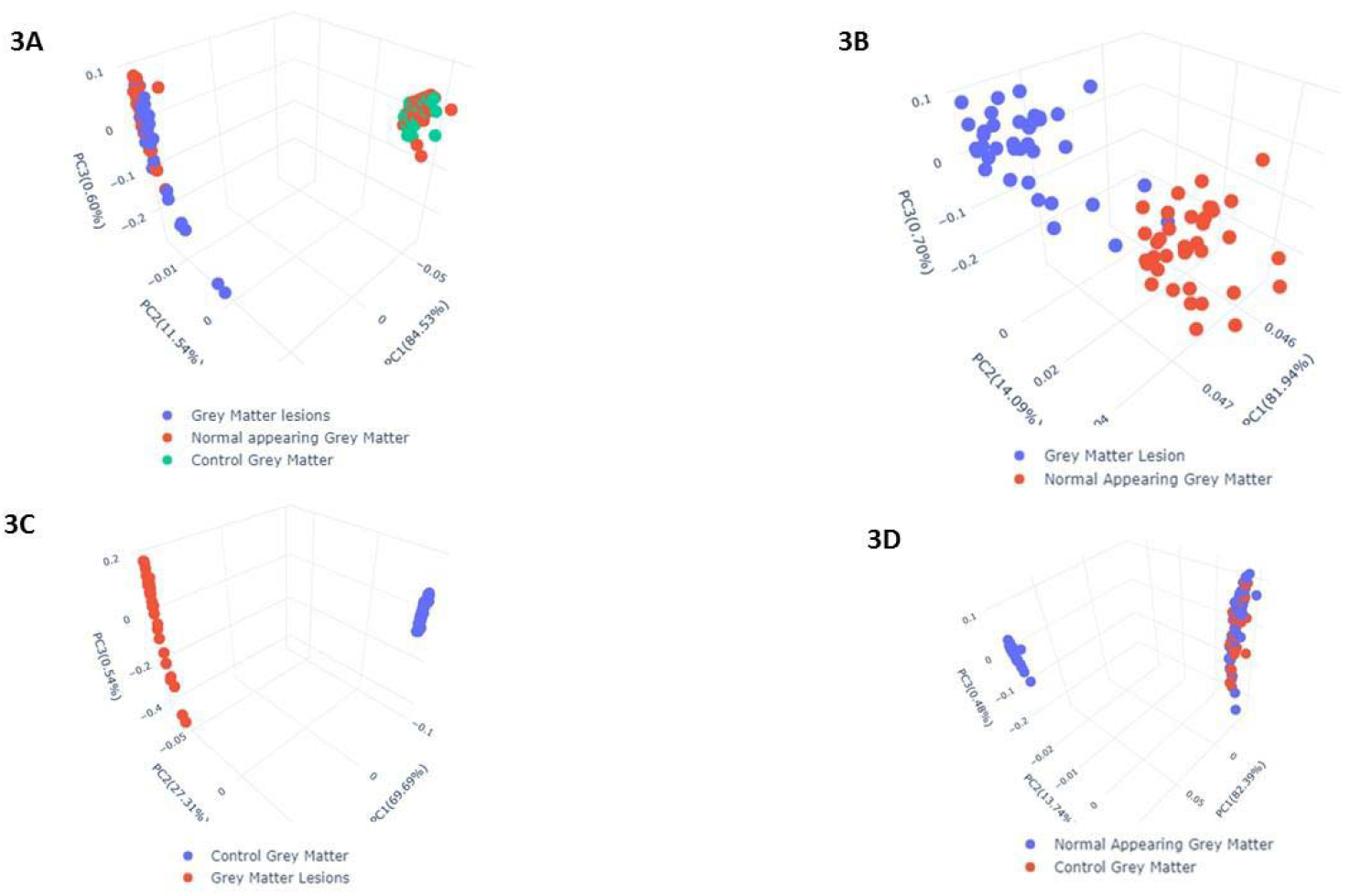
(A) PCA for All 3 conditions (Grey matter lesions Vs Normal appearing Grey Matter Vs Healthy Control Samples), (B) PCA For Grey matter lesions Vs Normal appearing Grey Matter, (C) PCA showing Variance among Control Grey matter(Healthy Control) and Grey Matter Lesion, and (D) PCA For Normal appearing Grey matter Vs Control Grey Matter samples.

### Downstream Analysis

It is evident from the PCA results (**Figure 3A to 3D**) that the variation between Grey Matter lesions and Control grey matter samples is maximum. Hence, we further carried out a Comparative analysis between Grey matter lesions and Healthy Control samples.Based on Figure 3C, it can be seen that healthy[Control] samples and the Grey matter lesion samples are well separated, which means there is significant variation between these groups. Hence, in downstream analysis, we performed differential gene expression analysis between Grey Matter lesions and Healthy control samples. When the Grey Matter lesion samples and Control samples were examined in a specific scatter plot, it was obvious that there was an improvement in principal components, and the two groups were separated from each other. After the observed variation between the these two groups-Grey Matter lesions and Control samples, the reason for this difference and its impacts could be investigated at the level of gene regulation.

### Differential Genes Expression Analysis

The differential gene expression analysis between the Grey matter lesions (GML) and Control grey matter (CGM) samples scrutinized 1525 significantly (p.adj value <0.05, Fold change >= ±1.5). Among them, 438 genes were found to be significantly upregulated (p.adj value <0.05, Fold change >= +1.5) genes and 1087 genes were found to be significantly downregulated (p.adj value <0.05, Fold change <= −1.5) in GML in comparison to CGM.

### Clustering and Heat Map revealed variations among Grey Matter lesions and Control samples

Hierarchical Clustering (26) was performed to understand whether significant genes are capable to form distinct clusters of Grey Matter lesions and Control Grey Matter samples based on their gene expression. Clustering results clearly show the distinct clusters of Grey Matter lesions and Control Grey Matter samples as shown in Figure 4. Heatmap representing the expression pattern of significant genes among Grey Matter lesions and Control Grey Matter samples, as shown in Figure 5.

**Figure 4:**
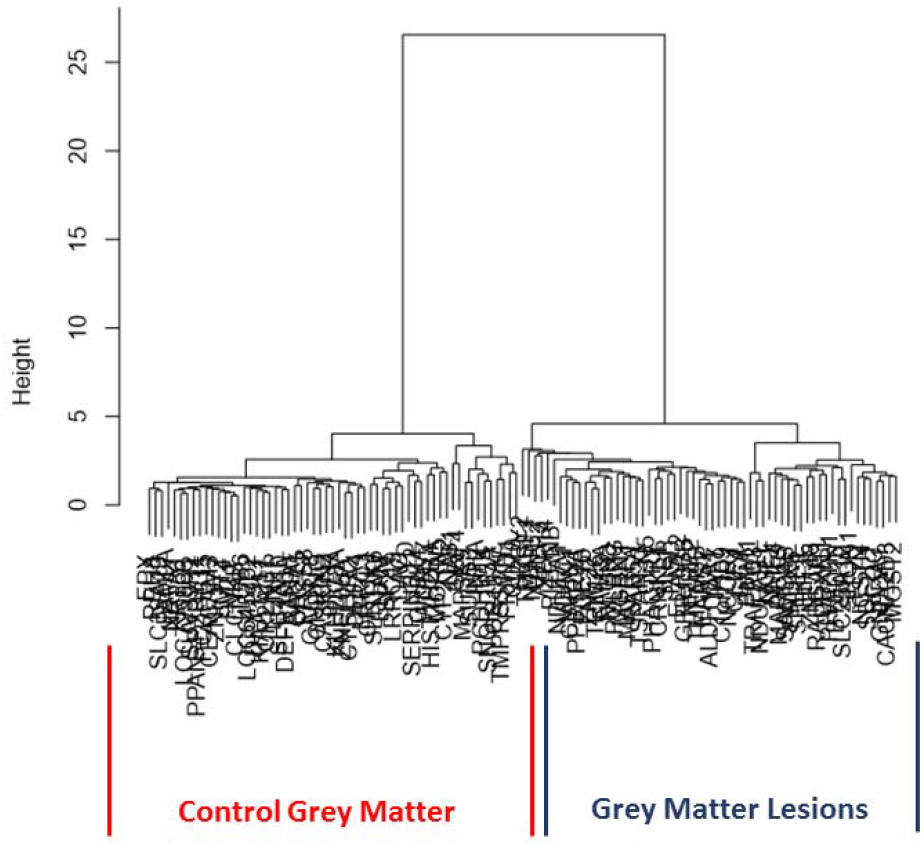
Hierarchical Clustering results as dendrograms. Red boxes clusters indicate the Control (Healthy) samples, and the blue one represents the clusters of the GML (lesions) samples.

**Figure 5:**
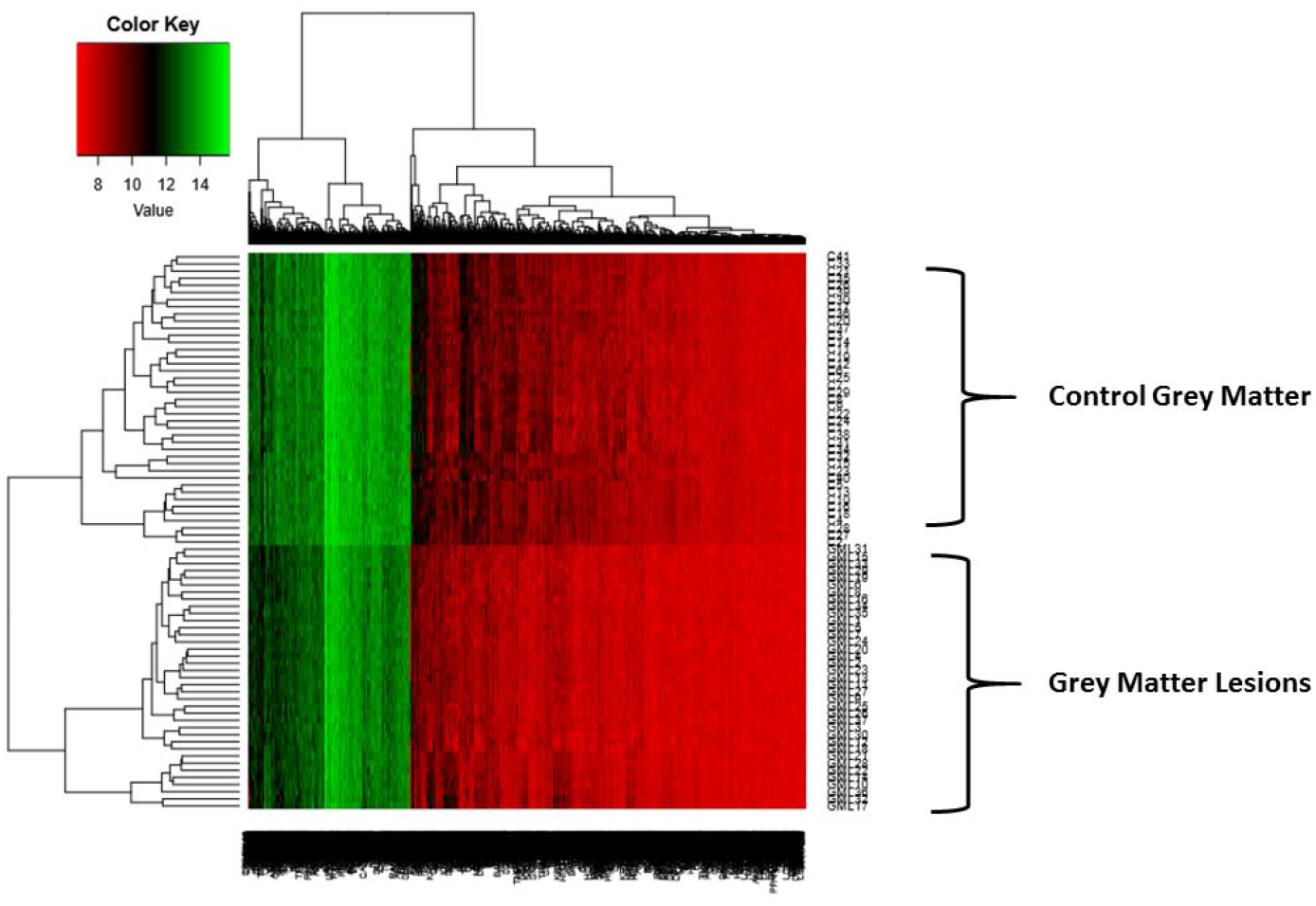
Heatmap representing the Gene expression patterns of significantly differentially expressed genes.

### Pathways involved in the pathogenesis of Multiple sclerosis

To understand the biological importance of pathways, gene ontology analysis was performed. Interestingly, although there were different pathways in upregulated genes, many represented biological pathways involved Huntington’s Disease, GTP Binding biological processes, Mitochondrial Chain 1 respiratory complexes (Figure 6A).

**Fig 6A:**
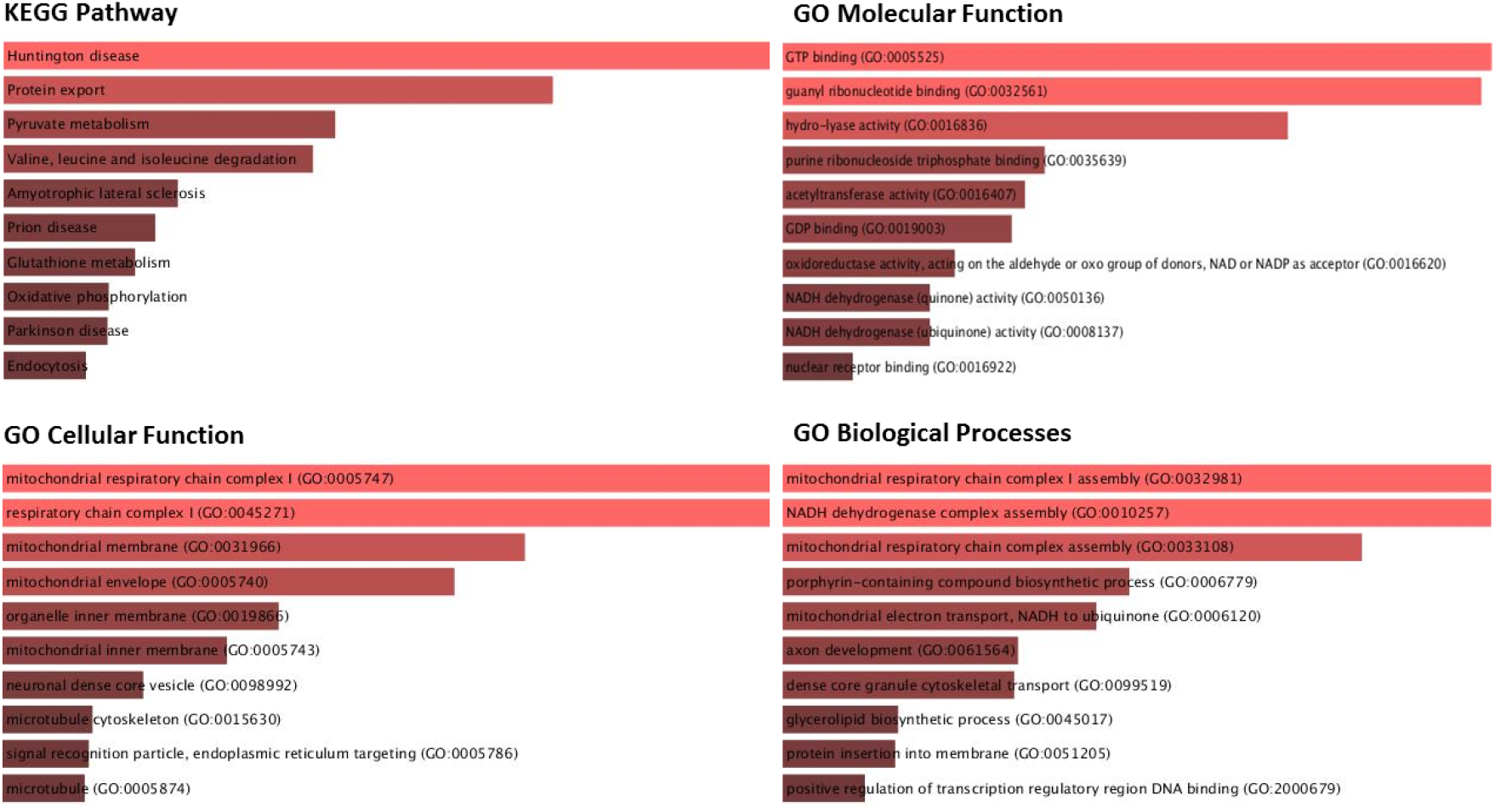
Gene ontology and KEGG pathway analysis of Up-regulated genes.

The same concept could be detected with down-regulated genes as shown in Figure 6B. The gene ontology analysis for down-regulated pathways involved Olfactory transduction pathways, Plasma membrane functional pathways, and the development of male primary sexual characteristics and gonad developmental pathways. According to the first principal component, Grey matter lesion had a lot of variability within its group (Figure 3A). Per the experiments conducted by Subbiah-Pugazhenthi(27), it shows that Cyclic AMP response element-binding protein (*CREB*), is a nuclear transcription factor that plays a major role in neurodegenerative diseases and it’s progression(27). Also, *CREB* is responsible for the retention of memory and survival of neurons (28). The role of the *TAF4* gene is increasing in neurodegeneration (29). Hence after a literature review on the close association of Huntington’s Disease (HD) associated genes involved/found in MS patients, we concluded. From our list of significantly expressed genes, many of the HD-associated genes were expressed differentially, specifically up-regulated. 7 HD-associated pathway genes that are significantly differentially expressed (up-regulated) in GML samples with their annotations are enlisted in Table 2.

**Fig 6B:**
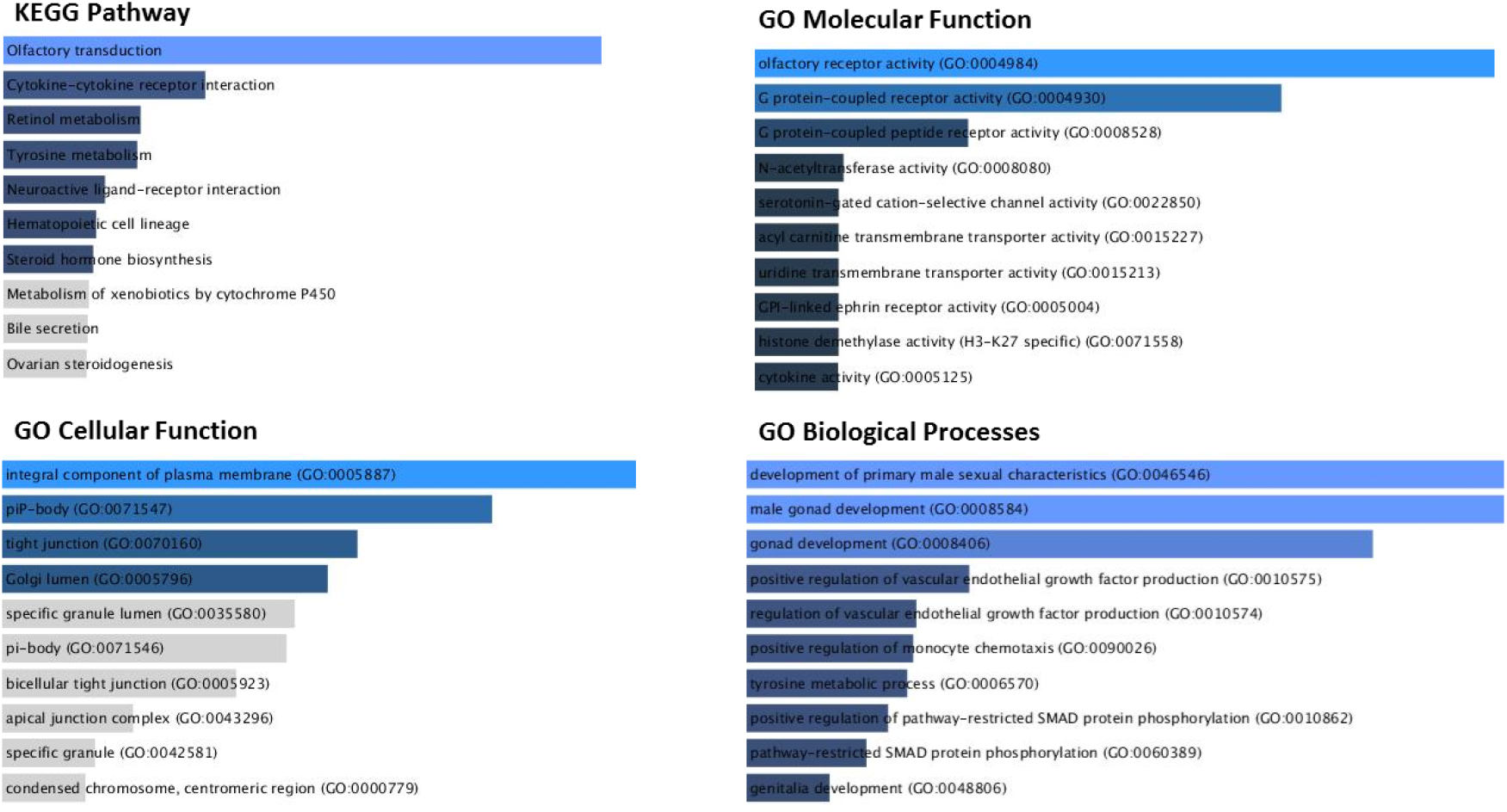
Gene ontology and KEGG pathway analysis of Down-regulated genes.

**Table 2.**
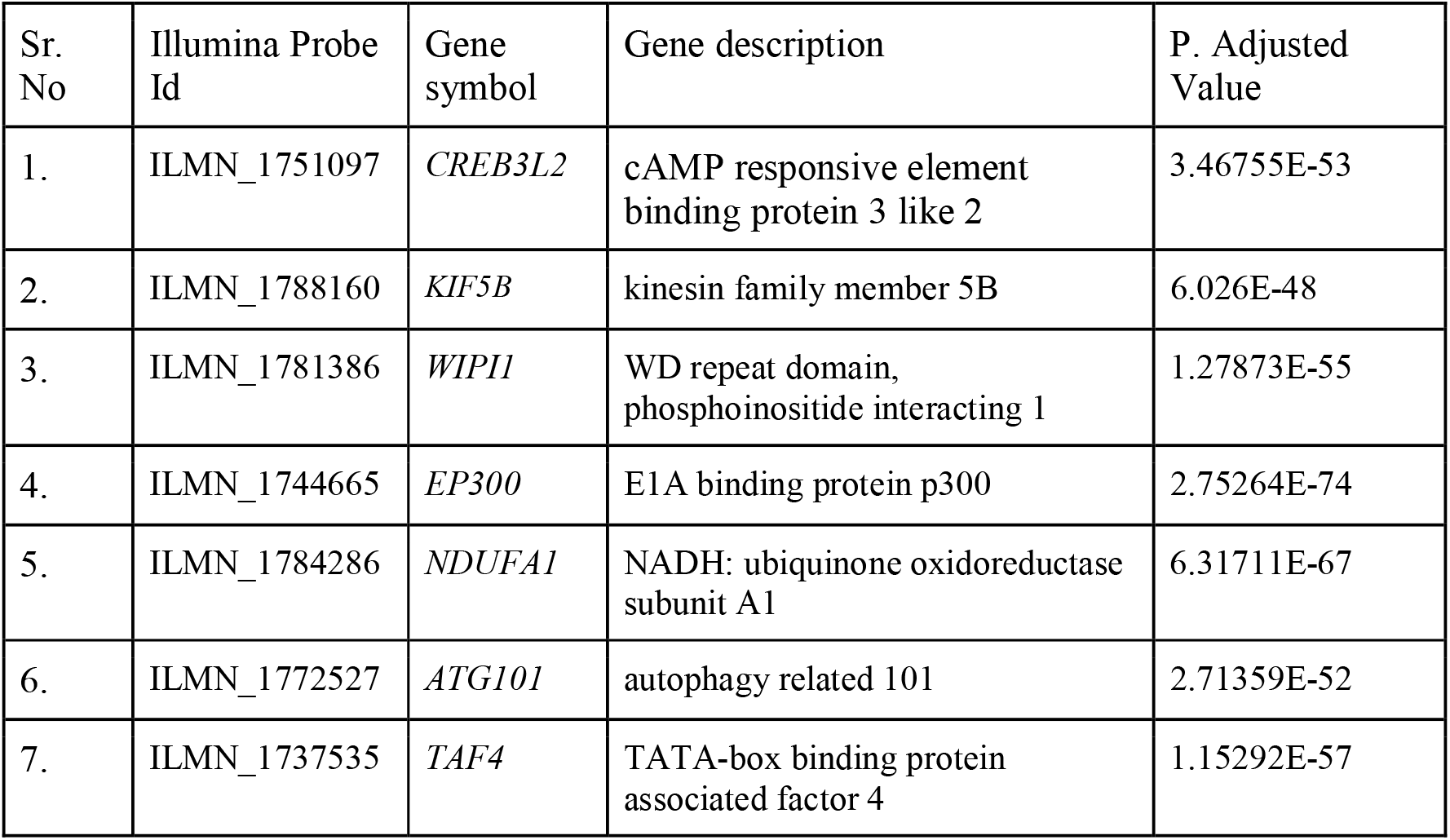
Differentially Expressed 7 Huntington Disease-associated Genes identified in our study (P. Adjusted value < 0.05;Fold change(>= ±1.5).

We can see in the heat map (Supplementary Figure S1) that the majority of Huntington’s associated genes are up-regulated in patients with multiple sclerosis lesions in comparison to the control samples. This was evident even in KEGG analysis (Figure 6A) as the Huntington Disease-associated pathway is significantly Up-regulated in Grey matter lesions (MS).

### Validation of the Results

Next, to assess and validate the 7 Huntington’s disease-associated genes expression patterns, we analyze the expression pattern of those 7 Huntington’s disease-associated genes in a validation dataset containing patients of MS without lesions. It is evident from the Heat-map (Supplementary Figure S1) that the 7 Huntington’s disease-associated genes in Multiple sclerosis normal-appearing Grey Matter are highly upregulated in comparison to control samples in the discovery dataset. Hence, we also analyze the expression pattern of these genes in the validation data, the bar-plot (Supplementary Figure S2) between Healthy (controls) and Normal appearing Grey matter without lesions (Multiple sclerosis) represents the expression pattern of the genes. Our validation results confirm our hypothesis. In the PCA plot (Supplementary Figure S3), we can clearly see the distinct clusters. Similarly, hierarchical clustering results (Supplementary Figure S4) confirming that the two groups form distinct clusters separately in the dendrogram, which implies these significant genes might predict MS characteristics.

### Discussion & Conclusion

The origin of Lesions in Multiple Sclerotic patients remains unknown. Previous studies addressing this question have shown contrasting results, thus making it difficult to establish any one of the theories as to the answer (30). In this study, we have attempted to reveal the close association of Multiple sclerosis with several affected Biological pathways namely Huntington disease-associated pathways using RNA-Seq Analysis. To identify differences in gene expression between Grey Matter lesion samples and Healthy control samples, RNA-Seq analysis was performed on RNA-seq samples obtained from both of the samples. Exploratory data analysis using PCA revealed that the samples of both categories form separate clusters, depicting the vast genetic differences between the MS Grey matter lesions and Control samples. This hints towards the theory that there is something that triggers the progression of MS diseases which leads to the formation of lesions at the transcriptomics level. Further studies on the same would be able to divulge the series of events that take place during this transformation.

Based on differential gene expression analysis using Welch’s T-Test, we observed 438 genes were significantly (adjusted P-value <0.05, Fold change (>= ±1.5) upregulated and 1087 (adjusted P-value <0.05, Fold change (>= ±1.5) genes were downregulated in GML in comparison to CGM. Next, the PCA plot and dendrogram from Hierarchical clustering indicated their significance in distinguishing GML and CGM. A heat map generated from these genes showed that most of the genes were downregulated in GML samples whereas some of them were up-regulated in GML samples, highlighting differential expression in both populations.

Gene ontology analysis performed using Enrichr based on significant gene sets showed obvious involvement of genes, specifically in Grey matter degeneration, via functional annotation and clustering. However, hits were obtained for genes associated with cell adhesion, Huntington’s Disease, Mitochondrial respiratory chain complexes, GTP-Protein synthesis, Olfactory and recovery adaptation pathways, migration, and invasion, indicating that their possible dysregulation may have led to the formation of Grey Matter lesions. Further, pathway analysis demonstrated a better understanding of the involvement of the gene sets in pathways representing the Odour dysfunction in MS patients which is based on downregulation of Olfactory transduction pathways. It also shows the up-regulation of pathways like Huntington disease-associated pathways, Mitochondrial respiratory chain complex pathways, and GTP Protein binding pathways. This study shows that as per our KEGG Analysis pathways, the most affected pathway is the HD-associated pathway. The associated genes with this pathway are *CREB3L2*, *KIF5B*, *WIPI1*, *EP300*, *NDUFA1*, *ATG101*, *AND TAF4*[as per Enrichr]as the key features that may substantially contribute to loss of cognitive functions in neurodegenerative disorders like Multiple sclerosis, Huntington’s disease, and further on. However, since our study was focused on samples of patients with multiple sclerosis, the involvement of certain genes that are also associated with HD is something of great importance. Studies have shown that the CREB Factor plays a major role in memory formation and neuronal regeneration (31). The gene *TAF4* plays a significant role in controlling the differentiation of human neural progenitor cells (32). Since *TAF4* is up-regulated in MS patients, in this study we have drawn a possible association between *TAF4* and neuronal degradation (33). *TAF4* is primarily concerned with the production of human neural progenitor cells. However, from our results depicted here, due to the significant up-regulation of *TAF4*, the production of new neural cells especially glial cells and neurons is hindered-leading to progressive neuroinflammation and neurodegeneration. Our results also depict, severe up-regulation of the *CREB* gene which might be the possible causes of progressive neuronal degeneration and loss of cognitive functions in patients of MS. The indication of such genes up-regulated in MS patients could be used as a predictive tool for identification that MS is progressing and this progression might be fatal and lead to severe other neurodegenerative disorders. Talking about the *KIF5B* gene, it is a motor neuron protein that plays a significant role in dendritic transport, synaptic plasticity, and memory retention (35). *WIPI1*(WD Repeat Domain, Phosphoinositide Interacting 1) is a proteincoding gene responsible for neurodegeneration because of its involvement in brain iron accumulation. Enhanced iron uptake in acute cases of MS has been reported which shows that the *WIPI1* gene can play a leading cause in iron accumulation and neurodegeneration of the brain (36).

It should be also noted *WIPI1*, *TAF4*, and *CREB* protein-coding genes are the ones that are also associated with the early onset of Huntington’s disease (37). Several studies have suggested that impairments of the autophagic process are associated with several neurodegenerative diseases like HD and MS (38). The depiction of these in patients of GML suggests that they are early predictive biomarkers for HD. Thus, while assessing the patients with MS, the Up-regulation of these genes can be used as a tool for recognizing progressive fatal neurodegeneration and onset of HD.

However, it should be noted that in this study we have portrayed 7 genes (*CREB3L2*, *KIF5B*, *WIPI1*, *EP300*, *NDUFA1*, *ATG101*, *AND TAF4*) as the key features that may substantially contribute to loss of cognitive functions and progressive neurodegeneration involved in the MS pathogenesis. This study was able to identify several protein-coding genes whose role in the association with HD remains largely unclear. Studying their functionality in the future may reveal key biomarkers or drug targets that could be exploited for MS in association with HD. People suffering from MS who might be potentially screened for HD can be looked upon for these genes in order to get signals for early onset of HD.

Conclusively, our study revealed significant differences in gene expression between Grey matter lesions and Healthy Control samples. Importantly, we confirm and validate the significant upregulation of 7 HD pathway-associated genes in GML in comparison to normal samples. This most likely reveals an important clue to the etiology of this fatal neurodegenerative disease.

### Future Directions

In the future, intensive research is required to assess the precise effect of Multiple sclerosis on genes like *NDUAF1*, *EP300*, *and ATG101*. More comprehensive research is needed to support these findings stated above. As part of a future study, it would be interesting to understand the changes in MS pathogenesis and its effects on olfactory transduction, Development of Primary male sexual characteristics, GTP Binding protein, and Plasma Membrane component pathways.

## Author Contributions

Rutvi Vaja proposed the research idea, performed the data analysis, and produced the figures and manuscript. Dr. Harpreet Kaur reviewed the data, provided assistance with analysis and interpretation, and supervised the whole project. Elia Brodsky and Dr. Mohit Mazumder provided expert reviews of the project.

## Competing Interests

The authors declare no competing financial and non-financial interests.

## Supplementary Data

Supplementary File 1: Supplementary Figures.

Supplementary File 2: Supplementary Tables.

## Acknowledgment

This research was performed within the framework of the Research fellowship program and was supported by Pine Biotech. Data processing and visualization pipelines were run with the T-Bio info Server.

## Abbreviations

MS: Multiple Sclerosis
GML: Grey matter lesions
CGM: Control Grey matter
CG: Control Samples
NAMG: Normal appearing Grey Matter
CNS: Central nervous system
HD: Huntington’s Disease
WIPI1: WD Repeat Domain, Phosphoinositide Interacting 1
CREB: cAMP response element-binding protein

## Notes

### Competing Interest Statement

The authors have declared no competing interest.

