## Supplementary File 1 for "In silico Analysis of Transcriptomic Profiling and Affected Biological pathways in Multiple Sclerosis"

**Emails of Authors:**

**Correspondence:**

*Rutvi Vaja

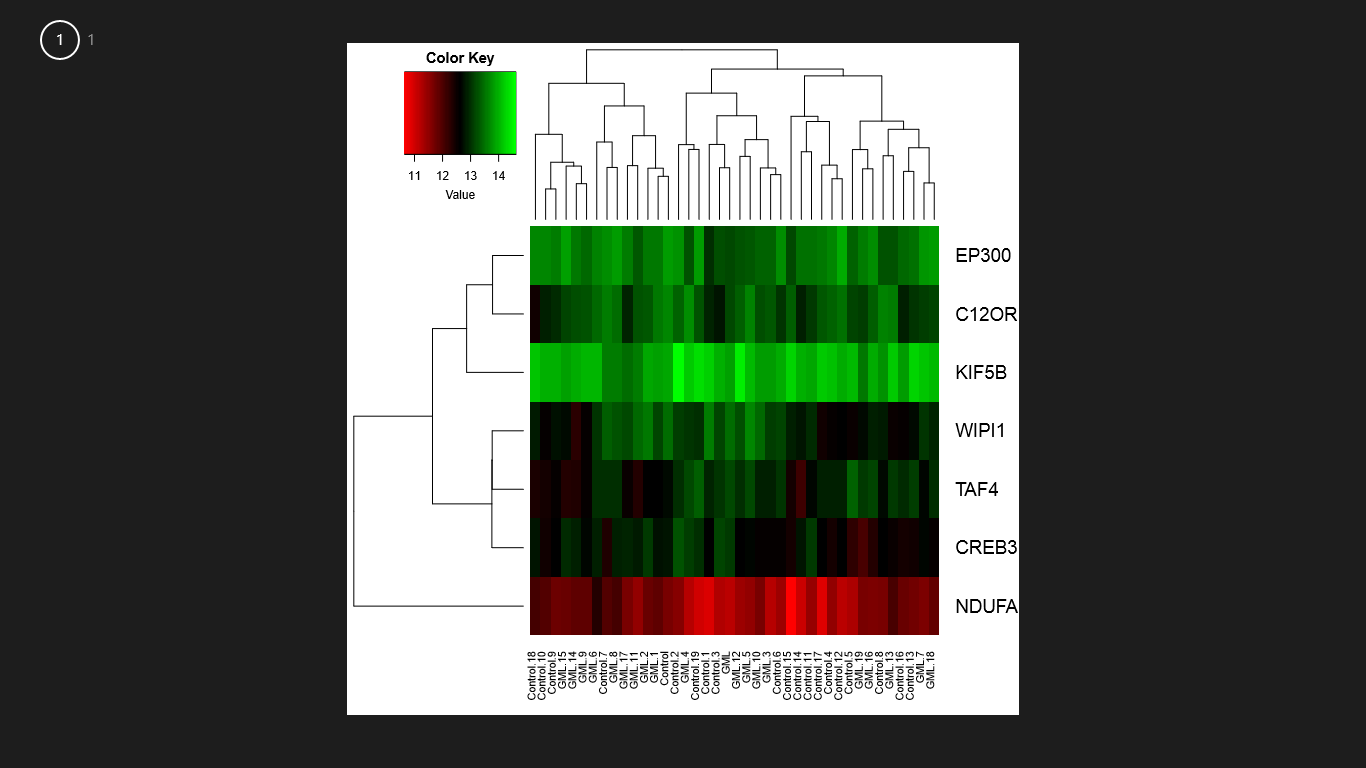

**Supplementary Figure S1: Heat Map representing the expression levels of Huntington Disease associated genes in Multiple sclerosis as well as Control (healthy) patients of the Discovery Data Set.**

BAR PLOT- HD Associated genes

|  | CREB3L2 | KIF5B | WIPI1 | EP300 | NDUFA13 | C12ORF44/ATG101 | TAF4 |
| --- | --- | --- | --- | --- | --- | --- | --- |
| NAGM | 13.0884 | 13.8938 | 13.5047 | 13.1972 | 11.1828 | 13.2156 | 13.1881 |
| NAGM | 12.7667 | 13.8974 | 13.1775 | 13.5516 | 11.9108 | 13.5715 | 12.6114 |
| NAGM | 13.106 | 13.9148 | 13.5742 | 13.5484 | 11.822 | 13.3222 | 12.629 |
| NAGM | 12.5559 | 13.8425 | 13.1221 | 13.3988 | 11.2227 | 13.3435 | 12.8739 |
| NAGM | 13.1178 | 14.211 | 13.0439 | 13.276 | 11.2328 | 13.7367 | 13.2247 |
| NAGM | 12.6764 | 14.071 | 13.6991 | 13.3446 | 11.5159 | 13.6616 | 13.2143 |
| NAGM | 12.8965 | 14.0509 | 13.0321 | 13.6626 | 12.3317 | 13.454 | 13.0286 |
| NAGM | 12.6936 | 14.1525 | 13.0639 | 13.7829 | 11.6584 | 13.1234 | 12.7019 |
| NAGM | 12.8904 | 13.5924 | 13.3022 | 13.8538 | 12.1144 | 13.466 | 13.0283 |
| NAGM | 12.6981 | 14.0231 | 12.5706 | 13.4402 | 11.8804 | 13.2763 | 12.557 |
| NAGM | 12.5639 | 13.8343 | 13.4445 | 13.4037 | 11.7022 | 13.233 | 12.901 |
| NAGM | 12.8685 | 13.6129 | 13.454 | 13.3182 | 11.5165 | 13.2926 | 12.32 |
| NAGM | 12.6283 | 14.4965 | 13.2658 | 13.3015 | 11.4027 | 13.3694 | 12.9954 |
| NAGM | 12.5521 | 14.2001 | 12.5479 | 13.3028 | 12.0478 | 13.6035 | 13.1015 |
| NAGM | 12.8941 | 13.9634 | 12.3029 | 13.5683 | 11.8794 | 13.2356 | 12.3682 |
| NAGM | 12.9627 | 13.8727 | 12.7183 | 13.8701 | 11.8063 | 13.1644 | 12.3226 |
| NAGM | 12.3322 | 13.9502 | 12.8878 | 13.7302 | 11.6696 | 13.3834 | 13.1507 |
| NAGM | 12.9391 | 13.4731 | 13.189 | 13.6159 | 11.6872 | 12.9421 | 12.5277 |
| NAGM | 12.5797 | 14.0808 | 12.9226 | 13.8329 | 11.8554 | 13.1855 | 13.0037 |
| NAGM | 12.0384 | 13.5805 | 12.7228 | 13.5995 | 11.7004 | 13.1364 | 13.0744 |
| Control | 8.63488 | 8.24391 | 7.66859 | 7.70072 | 7.90337 | 9.06458 | 9.0719 |
| Control | 9.01141 | 8.59269 | 7.72383 | 7.49184 | 7.79692 | 9.16182 | 9.84366 |
| Control | 8.46059 | 8.42356 | 7.49386 | 7.9064 | 7.92779 | 9.24271 | 9.78383 |
| Control | 9.37079 | 9.11722 | 7.56409 | 8.04626 | 8.00133 | 10.0472 | 10.803 |
| Control | 9.87925 | 8.87394 | 7.6607 | 8.19339 | 8.27201 | 9.48547 | 9.52438 |
| Control | 8.70619 | 8.42245 | 7.50047 | 7.58187 | 7.71667 | 8.51219 | 8.28676 |
| Control | 8.85809 | 7.91354 | 7.44667 | 7.77685 | 7.83172 | 9.34217 | 8.90167 |
| Control | 7.80204 | 8.46005 | 7.39028 | 7.79596 | 7.62272 | 8.16166 | 9.09865 |
| Control | 8.62485 | 8.26052 | 7.56871 | 7.9845 | 7.49238 | 8.75823 | 8.69189 |
| Control | 9.01947 | 9.13344 | 7.63443 | 8.10589 | 8.27995 | 9.42047 | 9.43068 |

**Supplementary Figure S2: Bar Plot representing the expression of 7 Huntington Disease associated genes in Normal appearing Grey matter (without lesions) as well as Control (healthy) patient samples of the Validation Data set.**

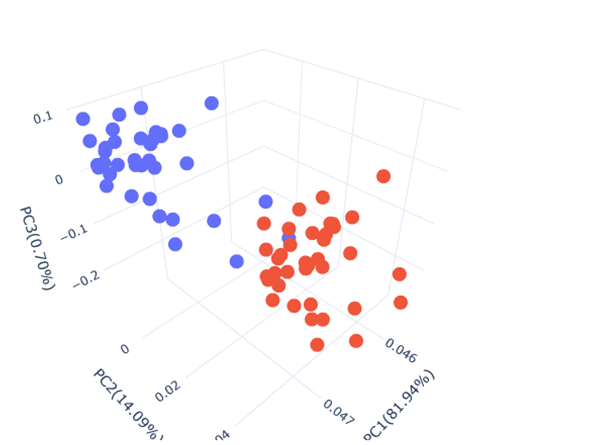

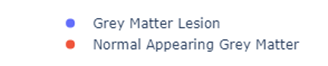

**Supplementary Figure S3: PCA based on significant genes showing Variance among Control Grey matter Healthy Control) and Normal appearing Grey Matter (Without Lesions) in the Validation data set.**

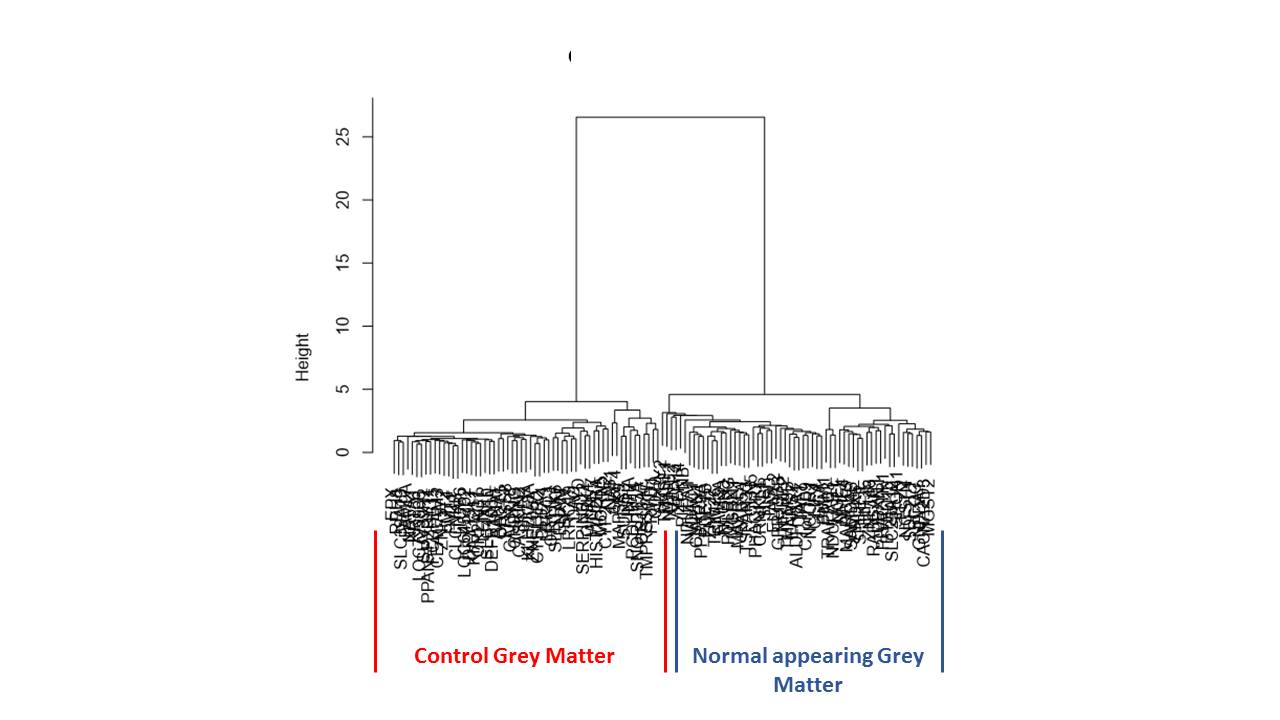

**Supplementary Figure S4: Hierarchical Clustering results based on Key genes in validation dataset. Each cluster in the dendrogram represents. Red boxes indicate the Control Healthy (highlighted in red colour) samples, and the NAGM (Normal appearing Grey Matter) samples highlighted in blue colour.**
